# Structure of the Cytoplasmic Ring of the *Xenopus laevis* Nuclear Pore Complex

**DOI:** 10.1101/2020.03.27.009407

**Authors:** Gaoxingyu Huang, Yanqing Zhang, Xuechen Zhu, Chao Zeng, Qifan Wang, Qiang Zhou, Qinghua Tao, Minhao Liu, Jianlin Lei, Chuangye Yan, Yigong Shi

## Abstract

Nuclear pore complex (NPC) exhibits structural plasticity and has only been characterized at local resolutions of up to 15 Å for the cytoplasmic ring (CR). Here we present a single-particle cryo-electron microscopy (cryo-EM) structure of the CR from *Xenopus laevis* NPC at average resolutions of 5.5-7.9 Å, with local resolutions reaching 4.5 Å. Improved resolutions allow identification and placement of secondary structural elements in the majority of the CR components. The two Y complexes in each CR subunit interact with each other and associate with those from flanking subunits, forming a circular scaffold. Within each CR subunit, the Nup358-containing region wraps around the stems of both Y complexes, likely stabilizing the scaffold. Nup205 connects the short arms of the two Y complexes and associates with the stem of a neighbouring Y complex. The Nup214-containing region uses an extended coiled-coil to link Nup85 of the two Y complexes and protrudes into the axial pore of the NPC. These previously uncharacterized structural features reveal insights into NPC assembly.

## INTRODUCTION

NPC is the only route of bidirectional cargo transport between the cytoplasm and the nucleus in a eukaryotic cell. With an estimated molecular mass of about 110-125 MDa in higher eukaryotes, NPC is among the largest and most important cellular machineries^1–6^. Detailed structural information is a pre-requisite for mechanistic understanding of NPC function. Towards this goal, the quest to elucidate three-dimensional structure of an NPC began more than 25 years ago^7^. X-ray crystallography has been successfully applied to the individual components and subcomplexes of the NPC, resulting in more than 30 crystal structures of individual domains and sub-complexes in the cytoplasmic ring and its associated cytoplasmic filaments^3,5^. For the intact NPC, however, the only viable method of structural investigation is electron microscopy (EM), which has yielded a reconstruction of low resolution in 1993 using single particle approach^8^ and about 10 reconstructions of moderate resolutions in recent years using cryo-electron tomography (cryo-ET)^5,9^. In particular, general structural features of the NPC have been revealed in *Dictyostelium discoideum*^10,11^, *Homo sapiens* (*H. sapiens*)^12,13^, *Xenopus laevis* (*X. laevis*)^14^, *Saccharomyces cerevisiae* (*S. cerevisiae*)^15^, and *Chlamydomonas reinhardtii*^16^. The cryo-ET reconstruction allows docking of known X-ray structures^12,13,15,17–19^.

The NPC has a cylindrical appearance and resides at the site of fusion between the inner and outer nuclear membranes. Along the cylindrical axis from the cytoplasmic side, an NPC consists of cytoplasmic filaments (CF), a cytoplasmic ring (CR), an inner ring (IR), a nuclear ring (NR) and a nuclear basket (NB)^3,5,20^. The ring scaffolds associate with one another and with the membrane. Within the lumen of the nuclear envelope (NE), there is a circular scaffold known as the luminal ring (LR)^8,21^.

The NPC displays an approximate eight-fold symmetry. The unusually large size of the NPC, together with its conformational plasticity, represents a challenge for structural determination at improved resolutions, particularly by cryo-ET. To date, local resolution of up to 15 Å has been achieved for the CR subunit of the human NPC through sub-tomogram averaging (STA) by cryo-ET^13^. At this resolution, protein components are only differentiated at the domain level and X-ray structures can be rigidly docked into the reconstruction to generate composite coordinates^13,17–19^. In this study, we report the cryo-EM structure of the CR from *X. laevis* NPC through single particle analysis (SPA). Our reconstruction displays average resolutions of 5.5-7.9 Å, which shows the tubular density for individual α-helices and allow placement of secondary structural elements for most components of the CR.

## RESULTS

### Single particle cryo-electron microscopy

The NE of *X. laevis* oocytes were prepared as described^21^. Cryo-ET only allows a relatively low dose of electrons at each defined tilting angle, which may hamper the resolution due to decreased signal-to-noise ratio. To overcome this drawback, we took the cryo-EM SPA approach. The grids were prepared and imaged using a K2 Summit detector mounted on a Titan Krios microscope. Because the NE was planarly spread on the grid, a majority of the NPCs had their cylindrical axes roughly perpendicular to the sample grid, creating an orientation bias. To alleviate this problem, the micrographs were recorded with the grid tilting at angles of 0, 30, 45 and 55 degrees^22^ (Supplementary information, Fig. S1). 12,399 good micrographs allowed manual picking of 289,306 NPC particles (Supplementary information, Table S1).

A subset of the data was used to generate an initial reconstruction of the CR at 18.3 Å resolution using a pixel size of 8.88 Å. This reconstruction already includes the known features of the CR^13,14^ (Supplementary information, Fig. S2) and allows generation of a preliminary reconstruction for the CR subunit at 9.4 Å resolution (Supplementary information, Fig. S3a). Application of this data processing strategy to the complete data set yielded a reconstruction of the CR at 17.8 Å resolution with a pixel size of 8.88 Å and the CR subunit at an average resolution of 8.0 Å with a pixel size of 2.22 Å (Supplementary information, Fig. S3b). To improve the local EM maps, three overlapping masks were applied to cover the CR subunit. The final average resolutions are 5.5 Å for the Core region, 7.1 Å for the Nup358-containing region, and 7.9 Å for the Nup214-containing region (Fig. 1a-c; Supplementary information, Fig. S3b). The particles for these regions exhibit good angular distributions (Supplementary information, Fig. S4a-c). The EM density maps are markedly improved compared to previous cryo-ET reconstructions^13^ (Supplementary information, Fig. S4d) and display features of secondary structural elements (Supplementary information, Figs. S5-9).

**Fig. 1.**
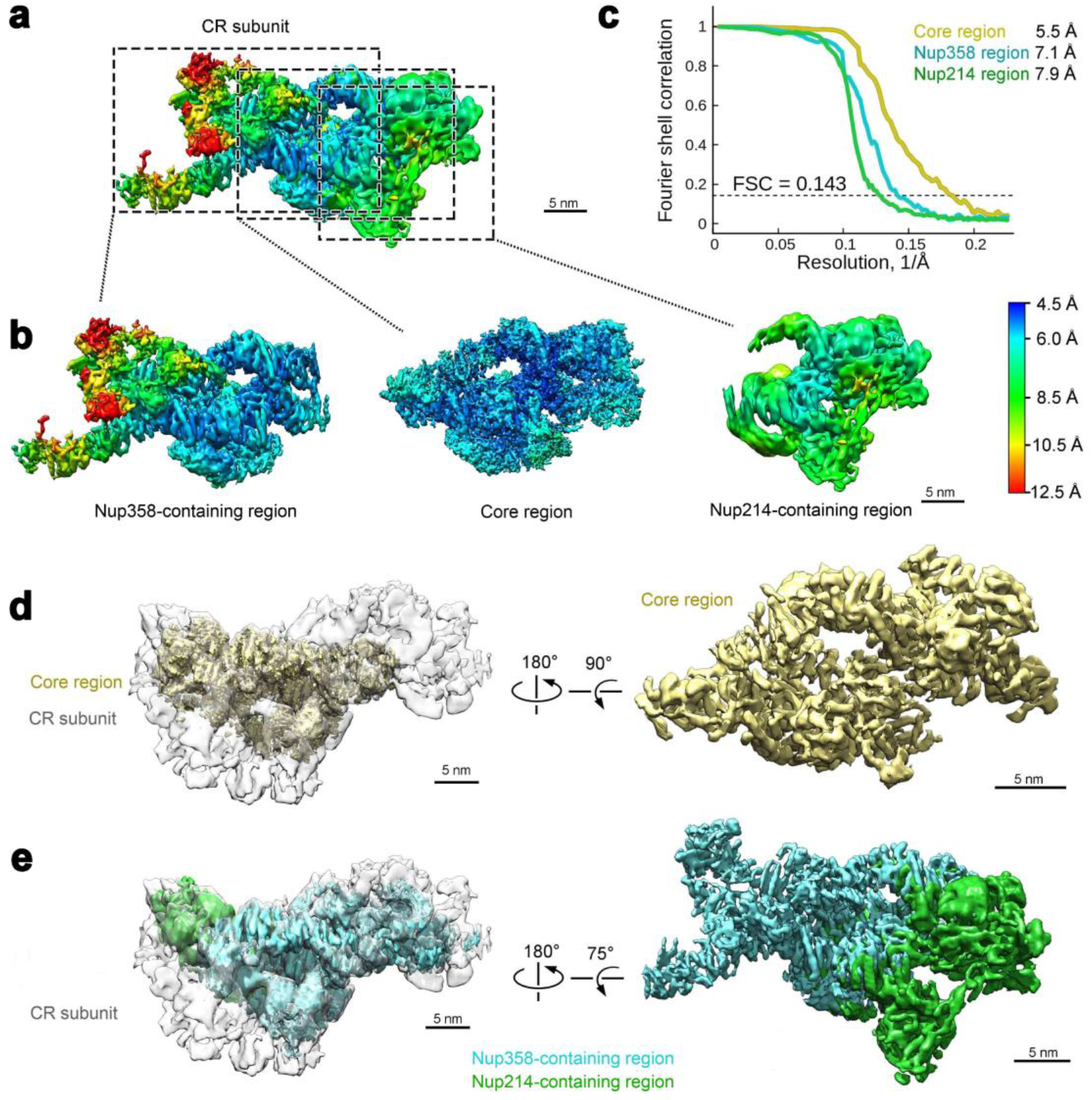
Cryo-EM structure of a subunit of the cytoplasmic ring (CR) of the *Xenopus laevis* (*X. laevis*) NPC. **a** The overall EM density map of a CR subunit. The local resolutions between 4.5 Å and 12.5 Å are color-coded. The CR consists of eight subunits. Each subunit is divided into three overlapping regions for soft mask application during data-processing: the Core region, the Nup358-containing region, and the Nup214-containing region. These three regions are indicated by dashed boxes. **b** The individual EM density maps for the Core, Nup358-containing, and Nup214-containing regions. The local resolutions are color-coded identically as in panel **a**. **c** The average resolutions for the Core, Nup358-containing, and Nup214-containing regions are 5.5 Å, 7.1 Å and 7.9 Å, respectively. Shown here are the Fourier shell correlation (FSC) plots of the reconstructions for the three regions. The resolutions were estimated on the basis of the FSC value of 0.143. **d** Cryo-EM reconstruction of the Core region of the CR subunit. Shown here is the EM density map of the Core region (colored yellow), which is placed in the CR subunit (left panel) and displayed in isolation (right panel). **e** Cryo-EM reconstruction of the Nup358-containing region and the Nup214-containing region. Shown here is the EM density map (colored cyan for the Nup358-containing region and green for the Nup214-containing region), which is placed in the CR subunit (left panel) and displayed in isolation (right panel).

### Overall structure of the CR

The average resolutions reach 4.5-5.5 Å for the bulk of the reconstruction in the Core region of the CR subunit, where densities for individual α-helices can be differentiated (Fig. 1b,d; Supplementary information, Fig. S5-9). The final reconstruction for the Nup358-containing region exhibits a wide range of local resolutions, from 5 Å in the center to 12 Å in one distant corner (Fig. 1b). Nonetheless, a proportion of the EM maps for the Nup358-containing region still displays tubular densities that are characteristic of α-helices (Fig. 1e; Supplementary information, Fig. S7). In contrast, the Nup214-containing region exhibits uniformly moderate resolution, mostly between 6 and 10 Å (Fig. 1b,e). The reconstructions for the three regions were merged into a single map for the CR subunit.

Eight copies of the CR subunit reconstruction were docked into the 17.8-Å map to generate a reconstruction of the CR (Fig. 2a, left panel). The CR measures 122 nm and 70 nm in outer and inner diameters, respectively. Composite coordinates have been previously obtained for the human Y complex^13,23^, which comprises a short arm (Nup85, Nup43, and Seh1), a long arm (Nup160 and Nup37), and a stem (Nup96, Sec13, Nup107, and Nup133). Using homology modeling, we generated composite coordinates of the *X. laevis* Y complex. Based on the EM maps, we manually adjusted individual secondary structural elements and orientation of the protein components. This effort was greatly facilitated by the characteristic EM density for about 200 α-helices, particularly those from Nup85 (Supplementary information, Fig. S5), Nup96, and Nup107. Notably, the assigned top and bottom faces of the Nup43 β-propeller (convention of Wall et al^24^) are opposite of those proposed for the human Nup43 (refs. 12,13). Two copies of the Y complex are present in each CR subunit (Fig. 2a, right panel) and are named inner and outer Y complexes relative to the center of the NPC (Fig. 2b). These two Y complexes occupy approximately 60 percent of the EM density map.

**Fig. 2.**
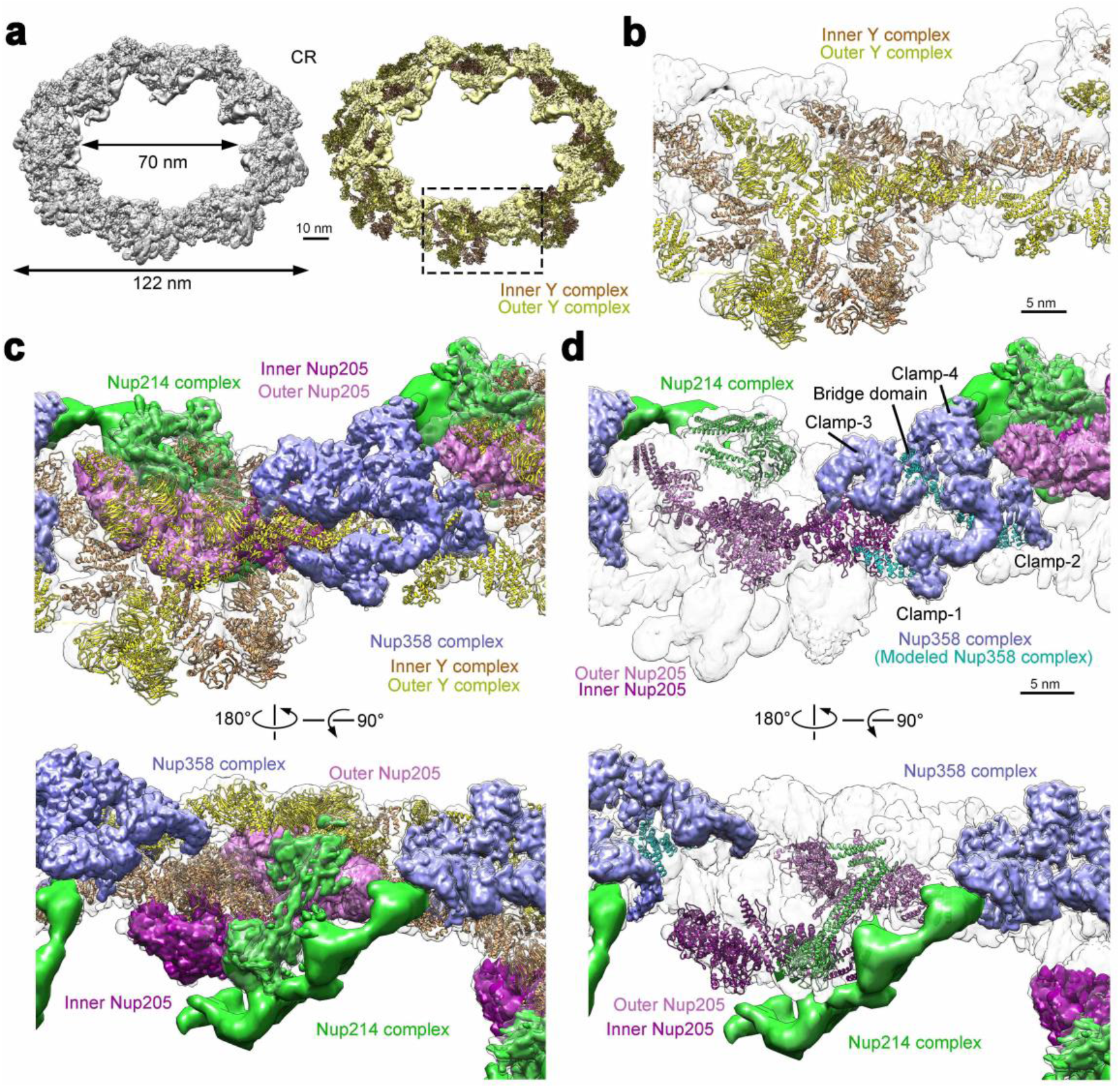
An atomic model of the CR from the *X. laevis* NPC. **a** Overall structure of the assembled CR at a tilting angle of 45°. Eight copies of the final reconstruction of the CR subunit were docked into the 17.8-Å EM density map of the CR to generate a complete structure (left panel). The coordinates of the *X. laevis* Y complex were generated through homology modeling and adjusted on the basis of the EM density map. Two copies of the Y complex (colored dark yellow and salmon) were fitted into the density map of each CR subunit. The EM density that cannot be accounted for by the Y complexes is shown and colored yellow (right panel). One CR subunit is boxed. **b** A close-up view on the two Y complexes in one CR subunit. Among the two Y complexes, the one located closer to the pore is named the inner Y complex (colored salmon) and the other is termed the outer Y complex (yellow). The EM density yet to be assigned is shown in grey. **c** The EM density maps of the Nup358 complex (colored purple), the Nup214 complex (green), and Nup205 (magenta) in each CR subunit. Two views are shown. **d** Placement of secondary structural elements in the Nup358 complex (colored cyan), the Nup214 complex (green), and Nup205 (magenta) in each CR subunit. Two views are shown. In addition to the deposited structures of the two Y complexes, the main chains of approximately 5,500 amino acids were placed into the EM density maps in our study.

The unoccupied EM density in each CR subunit is mainly distributed in three distinct regions. Decreased cellular expression of Nup358 and Nup214 results in marked and specific reduction of EM density in two of these three regions^12,13^; therefore, the proteins in these two regions are collectively referred to as the Nup358 complex and Nup214 complex (Fig. 2c). The third region had been previously assigned to Nup188 or Nup205 (refs. 12,13,17). Based on the EM maps, we placed 45 α-helices into the Nup358 complex, six α-helices and a β-propeller into the Nup214 complex, and 55 α-helices into two molecules of Nup205 or Nup188 (Fig. 2d; Supplementary information, Figs. S7 & 8). In addition, we modeled a hybrid helical domain (referred to as Nup96N/160C) that consists of seven α-helices from the N-terminal domain (NTD) of Nup96, five α-helices from the C-terminal domain (CTD) of Nup160 and one α-helix likely from Sec13 (Supplementary information, Fig. S6a). In addition to these 132 α-helices, the EM density also shows tubular features for about 120-150 α-helices that either merge into each other or are connected to neighbouring helices.

We have generated a main chain model of the CR subunit that includes about 16,000 amino acids. Among these residues, approximately 13,500 have been assigned to α-helices and β-propellers based on EM maps, which include 800 from the Nup358 complex, 780 from the Nup214 complex, and 3,550 from two molecules of Nup205 or Nup188. The EM density for the Nup358 complex comprises four characteristic Clamps^13^ and a newly identified bridge domain (Fig. 2d). The EM density for the Nup214 complex has a Y-shaped appearance (Fig. 2c, lower panel), of which one arm displays fine features and have been modeled into two coiled-coils and a β-propeller (Fig. 2d, lower panel). The EM density for Nup205 or Nup188 was assigned to two molecules of Nup205 (Fig. 2c,d), not Nup188 as previously proposed^12,17^.

### The Y complex as the scaffold of the CR

The improved resolution of the EM map allows accurate modeling of two Y complexes (Supplementary information, Fig. S10a). The inner and outer Y complexes within each subunit pair up in the same head-to-tail fashion; eight pairs of the Y complexes interact with each other to form a closed ring scaffold (Fig. 3a). Consistent with published results, the inner and outer Y complexes contact each other through two distinct interfaces within the same CR subunit (Supplementary information, Fig. S10b). At one interface, Sec13 and Nup96 at the vertex of the inner Y complex interact with Nup107 in the stem of the outer Y complex^13,19^ (Supplementary information, Fig. S10c). Nup133 consists of an N-terminal β-propeller followed by an extended α-helical domain^25,26^. At the other interface, the C-terminal α-helices of Nup133 from the inner Y complex (inner Nup133) interact with α-helices at the N-terminal end of the extended α-helical domain of Nup133 from the outer Y complex (outer Nup133) in the same subunit^13^ (Supplementary information, Fig. S10d).

**Fig. 3.**
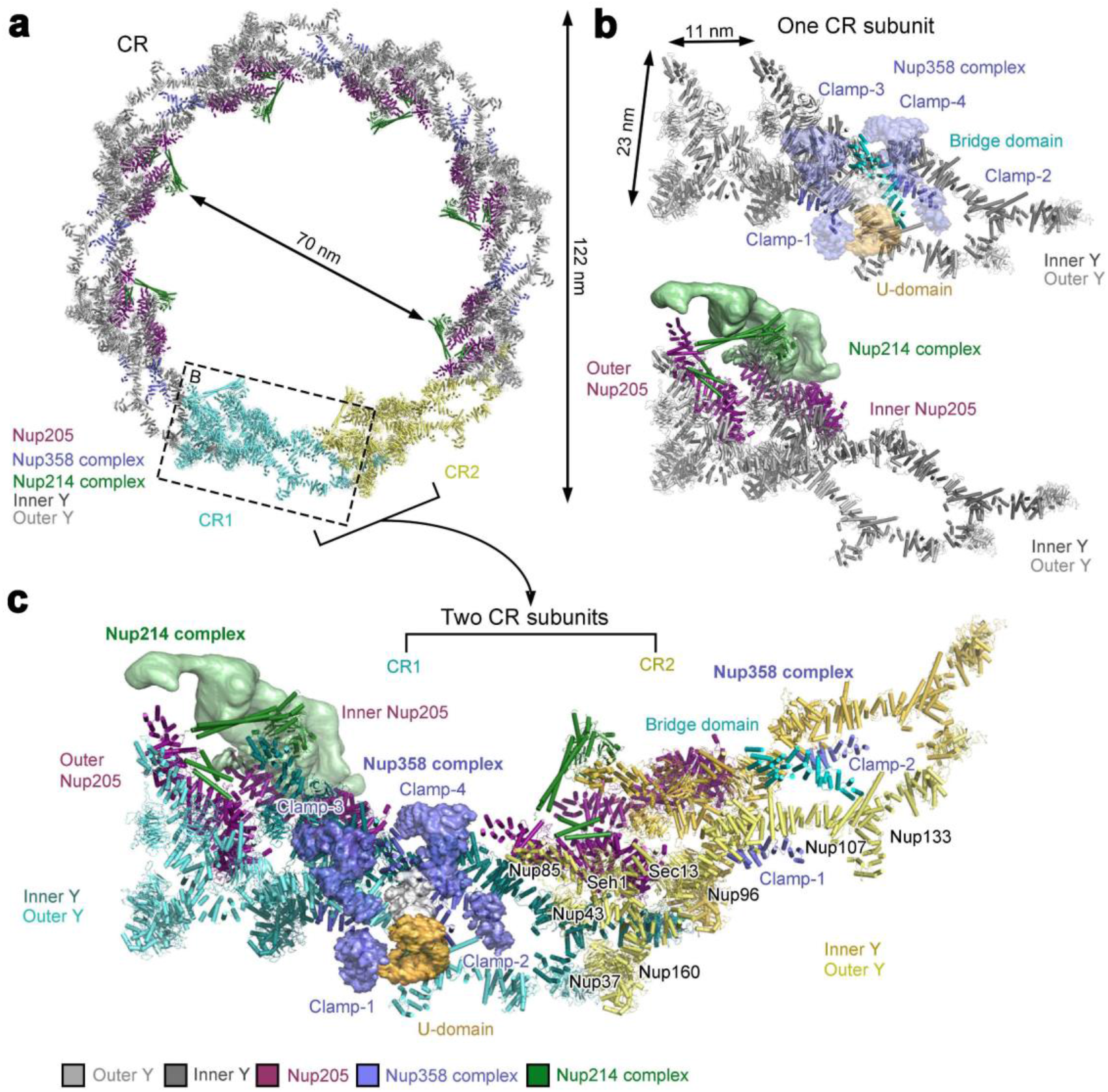
Overall organization of the CR. **a** Overall structure of the CR. The main-chain model of the CR contains approximately 108,000 amino acids, amounting to a combined molecular mass of nearly 11.9 MDa. Except for two subunits (colored cyan and yellow), subcomplexes in the other six subunits are color-coded. The Y complex, the Nup358 complex, the Nup214 complex, and Nup205 are colored grey, purple blue, green, and magenta, respectively. The 16 Y complexes assemble into a closed ring scaffold, on which the Nup358 complex, the Nup214 complex, and Nup205 associates. **b** Close-up views of a CR subunit. Within the same CR subunit, the inner and outer Y complexes associate with each other through two distinct interfaces: one between the inner vertex and outer stem and the other between the inner and outer stems (upper panel). The four Clamps of the Nup358 complex wrap around the stems of the two Y complexes. The two molecules of Nup205 directly interacts with the short and long arms of the two Y complexes (lower panel). The Nup214 complex interacts with the Y complex and Nup205. **c** A close-up view of the interface between two neighbouring CR subunits. The long arms of the inner and outer Y complexes directly associate with the stems of the inner and outer Y complexes from the neighbouring subunit, respectively. In addition, outer Nup205 binds the stem of the inner Y complex from the neighbouring subunit.

Between two neighbouring CR subunits, the N-terminal β-propeller of the inner or outer Nup133 associates with the N-terminal β-propeller and the ensuing α-helices of the inner or outer Nup160, respectively^13,17^ (Supplementary information, Fig. S10d). The extended α-helical domain of inner Nup133 also interacts with outer Nup37 and outer Nup160 of the adjacent subunit. Furthermore, the finger helix^27^ of inner Nup107 reaches out to contact the outer Nup43 and C-terminal α-helices of outer Nup85 from the adjacent subunit^13^ (Supplementary information, Fig. S10d). Together, these interactions allow the eight pairs of Y complexes to form a continuous circular scaffold.

Except the β-propeller of Nup133 at the distant tail, the inner and outer Y complexes exhibit a virtually identical conformation, with a root-mean-squared deviation (RMSD) of 5.6 Å over all main chain atoms (Supplementary information, Fig. S11a). Reflecting evolutionary conservation, the inner and outer Y complexes from *X. laevis* can be very well superimposed to those from *H. sapiens*^13^ (Supplementary information, Fig. S11b). More conformational variation is observed for the comparisons between the *X. laevis* Y complex and *Myceliophthora thermophila* (*M. thermophila*) Y complex^23^ or *S. cerevisiae* Y complex^19^. With the stem region aligned, the short arms of the *X. laevis* and *M. thermophila* Y complexes extend an angle of 40 degrees (Supplementary information, Fig. S11c). Despite a similar overall conformation between the *X. laevis* and *S. cerevisiae* Y complexes, there are many local differences (Supplementary information, Fig. S11d).

The Nup358 complex, the Nup214 complex, and Nup205 associate with the Y complexes (Fig. 3a,b). The four Clamps of the Nup358 complex wrap around the stems of the two Y complexes^13^ (Fig. 3b, upper panel). Clamp-1 and Clamp-3 sandwich the stem of the outer Y complex, whereas Clamp-2 and Clamp-4 fix the inner Y complex. The bridge domain closely associates with Clamps-3 and -4 and contacts the stem of the outer Y complex and the vertex of the inner Y complex (Fig. 3b,c). The two molecules of Nup205, dubbed inner and outer Nup205, closely interact with the short arms of the inner and outer Y complexes, respectively (Fig. 3b, lower panel). A prominent coiled-coil of the Nup214 complex connects the two short arms of the Y complexes, allowing the complex to project into the pore of the NPC (Fig. 3b,c). This coiled-coil likely corresponds to that formed by Nup214, Nup88, and Nup62 (refs. 28,29). In the assembled CR, Nup205 directly contacts the Nup214 complex within the same subunit, and the Nup214 complex from one subunit interacts with the Nup358 complex from the adjacent subunit (Fig. 3c).

### The Nup358 complex

As the largest protein component of the CR, Nup358 from *X. laevis* has an α-helical domain (residues 1-830) at its N-terminus^30–33^ (Fig. 4a). Eight Nup358 complexes are prominently positioned in the stem regions of the Y complexes (Fig. 4b). The assigned α-helices in each Nup358 complex include approximately 400 residues from the bridge domain and 200 each from Clamp-1 and Clamp-2 (Fig. 4c). For both Clamps, the 200 residues constitute 11 consecutive α-helices at one end of the Clamp and displays the same conformation (Fig. 4c; Supplementary information, Fig. S7b). In each case, α-helices 2–11 comprise five tetratricopeptide repeats (TPRs)^32^, with an extra α-helix placed at the N-terminus.

**Fig. 4.**
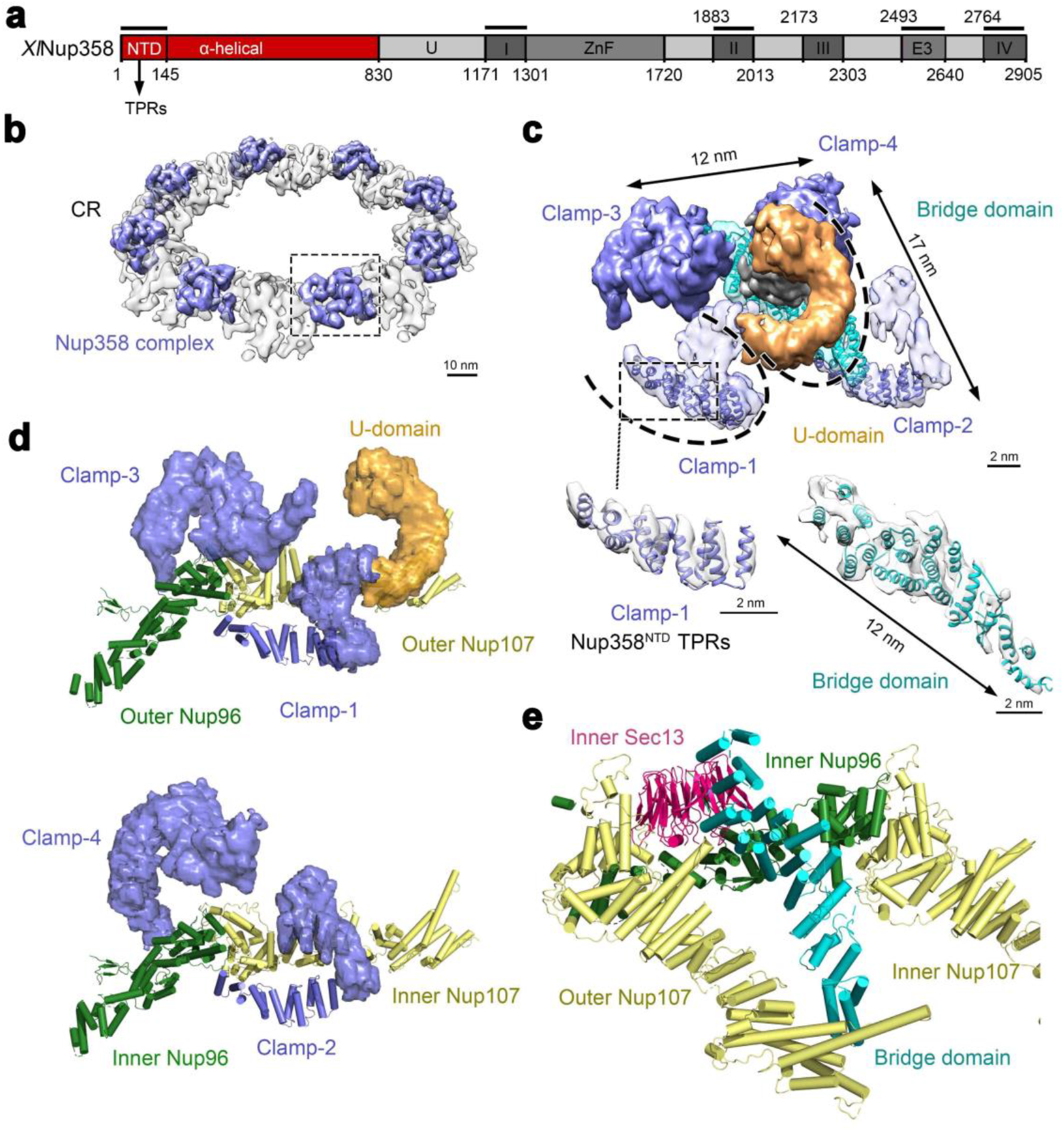
Four Clamps of the Nup358 complex sandwich the stems of the inner and outer Y complexes. **a** Domain organization of Nup358 from *X. laevis* (XlNup358). The various domains are drawn to scale. The thick black lines above the domain organization denote the fragments that have X-ray structures. In particular, the N-terminal domain (NTD, residue 1-145) of human Nup358 contains three TPRs (each encompassing a pair of α-helices)^32^. **b** The Nup358 complexes associate with the stems of the inner and outer Y complexes. Shown here is a cryo-EM reconstruction of the CR (transparent grey), with the Nup358 complexes shown in purple blue. **c** Structural features of the Nup358 complex. The Nup358 complex consists of four Clamps (Clamp-1, -2, -3, and -4, colored purple blue), and a bridge domain (cyan) in the center of the four Clamps. The ω-shaped Clamp-1 contains two U-domains, of which the second (orange) is out of plane with the first U-domain by nearly 90 degrees. The N-terminal 200-residue α-helices of Nup358 were modeled in Clamp-1 and -2. The bridge domain of 23 α-helices closely interacts with Clamp-4. The EM density maps for the N-terminal helices of Clamp-1 and the bridge domain are shown in the lower panels. **d** The Nup358 complex clamps the stem of the Y complex. Clamp-1 and Clamp-3 act like a pair of tweezers to hold the stem of the outer Y complex (upper panel). Similarly, Clamp-2 and Clamp-4 constitute a second pair of tweezers to hold the inner Y complex (lower panel). **e** The bridge domain of the Nup358 complex directly contacts inner Sec13, inner Nup96, and the finger helix of outer Nup107.

The X-ray structure for the NTD of human Nup358 (residues 1-145), which comprises eight α-helices and three TPRs^32^, can be fitted nicely into the EM density map for the N-terminal eight α-helices of Clamp-1 or Clamp-2. This analysis strongly suggests that Clamp-1 and Clamp-2 may represent the N-terminal sequences of Nup358. The features of the EM map for Clamp-3 and Clamp-4 are insufficient for placement of secondary structural elements. Nonetheless, the overall appearance of Clamp-3 or Clamp-4 is reminiscent of that for Clamp-1 or Clamp-2, suggesting the possibility that Clamp-3 and Clamp-4 may represent two additional copies of Nup358. This possibility is consistent with the proposal of four Nup358 molecules in each human CR subunit through quantitative mass spectrometry analysis^34^.

The continuous EM density for Clamp-1 constitutes an ω-shaped structure, which comprises two U-domains (Fig. 4c,d). The second U-domain is out of plane with the first by nearly 90 degrees (Fig. 4d, upper panel). In the low-pass filtered EM density map, Clamp-3 also displays an ω-shaped structure with two U-domains (Fig. 4b); in the final reconstruction, however, the second U-domain is no longer visible, likely due to its flexible nature. In contrast, only one U-domain has been unambiguously identified in Clamp-2 or Clamp-4 (Fig. 4d). The four Clamps do not directly interact with each other (Fig. 4c).

The Nup358 complex clamps onto the stems of both Y complexes within the same CR subunit (Fig. 4d). Clamps-1 and -3 act as a pair of tweezers to hold the stem of the outer Y complex (Fig. 4d, upper panel). The N-terminal α-helix of Clamp-1 directly contacts one side of outer Nup96; the two U-domains of Clamp-1 mainly interact with outer Nup107. Compared to Clamp-1, one end of the U-domain of Clamp-3 contacts the other side of outer Nup96 and the other end interacts with the junction between outer Nup96 and outer Nup107. In striking similarity to Clamps-1 and -3, Clamps-2 and -4 act as a second pair of tweezers to grip the stem of the inner Y complex (Fig. 4d, lower panel). In this case, inner Nup96 is sandwiched by the N-terminal α-helices of Clamp-2 and one end of Clamp-4.

The bridge domain resides in the center of the Nup358 complex and consists of 23 α-helices (Fig. 4c). It associates closely with Clamp-4 and to a lesser extent with Clamp-3. One end of the bridge domain directly binds the finger helix of outer Nup107 (Fig. 4e). The ridge of α-helices at the other end of the bridge domain contacts the lateral side of the β-propeller of inner Sec13 and the helical ridge of inner Nup96. The current resolution of our EM reconstruction is inadequate for sequence assignment of the bridge domain or determination of the helical directionality.

### Nup205 connects adjacent Y complexes

Nup205 is predicted to be almost entirely of α-helical conformation^35^. The *X. laevis* Nup205 comprises an NTD (residues 1-922), a middle domain (MID, residues 923-1391), and a TAIL domain (residues 1392-2011)^33,36^ (Fig. 5a). As the homologue of vertebrate Nup205, Nup192 from *Chaetomium thermophilum* (CtNup192) has been structurally characterized by X-ray crystallography^17,36,37^. In contrast to CtNup192 and *X. laevis* Nup188, *X. laevis* Nup205 contains extra 250 residues at the C-terminus (referred to as TAIL-C) that are predicted to be α-helices^38^. Two patches of S-shaped density in the cryo-ET reconstructions^12,13^ were thought to come from Nup188 on the basis of detectable cross-linking between Nup188 and Nup85 (refs. 12,17).

**Fig. 5.**
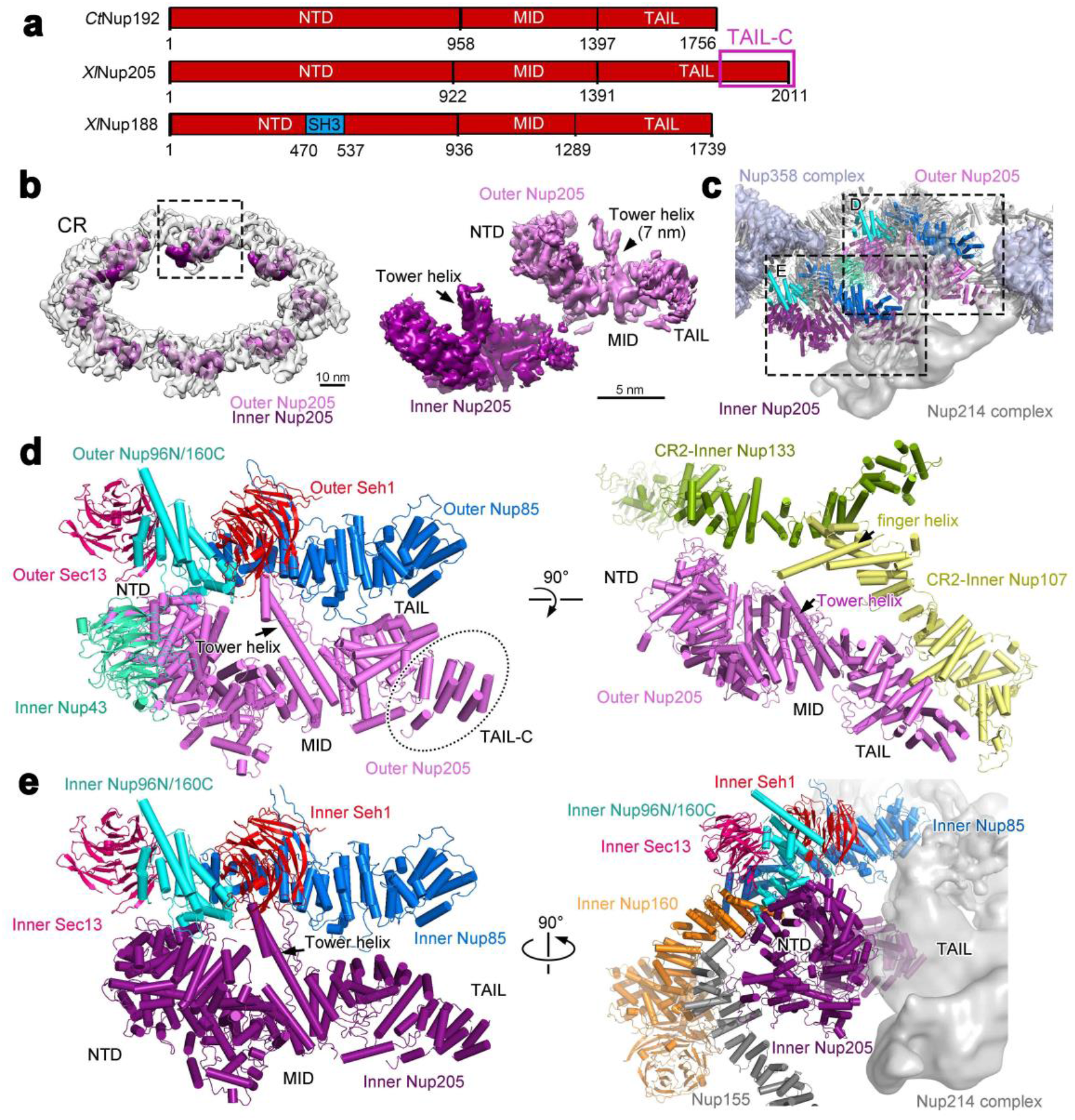
Nup205 interacts with the short arm of the Y complex and connects the Y complexes both within the same CR subunit and between adjacent subunits. **a** Domain organization of Nup205 and its comparison with Nup188 and Nup192. The domain organization of Nup205 from *X. laevis* (XlNup205) is displayed along with those of Nup188 from *X. laevis* (XlNup188) and Nup192 from *C. thermophilum* (CtNup192) for which the X-ray structure is available^17,36,37^. The various domains are drawn to scale. In particular, the C-terminal 250 amino acids of Nup205 (TAIL-C) are predicted to be α-helices but are not represented in the X-ray structure of the CtNup192 (ref. 17). **b** Nup205 directly interacts with the Y complex. The reconstruction of the CR is displayed in the left panel, with the EM density for Nup205 highlighted in magenta. The EM density maps for the two molecules of Nup205 are shown in the right panel. The Tower helix is clearly identifiable in both maps. **c** A close-up view on the locations of the two molecules of Nup205 in each CR subunit. Structures of the two molecules, named inner Nup205 (dark magenta) and outer Nup205 (magenta), are placed in the CR subunit. Proteins in the short arms of the two Y complexes are color-coded. The Nup358 complex is colored light purple blue. All other proteins are colored grey. The EM density map for the Nup214 complex is shown in transparent grey. **d** Outer Nup205 closely interacts with the short arm of the outer Y complex and the stem of the inner Y complex from an adjacent CR subunit. The NTD, Tower helix, and TAIL of outer Nup205 associate with the hybrid α-helical domain of outer Nup96N/160C, outer Nup85 and outer Seh1, and outer Nup85, respectively (left panel). The NTD also binds inner Nup43. Outer Nup205 also makes extensive contacts to inner Nup133 and inner Nup107 from the adjacent CR subunit (right panel). **e** Inner Nup205 binds the short arm of the inner Y complex (left panel) and the Nup214 complex (right panel). The interactions with the short arm are similar to those by outer Nup205 except that the TAIL of inner Nup205 no longer contacts inner Nup85.

In our final reconstruction of the CR subunit, such S-shaped density is present in the same location as that in the published cryo-ET reconstructions^12,13^. The S-shaped density in our reconstruction has considerably more features, with the Tower helices standing out (Fig. 5b). This density is highly unlikely to come from Nup188, because modeling of the full-length *M. thermophila* Nup188 (ref. 39) into the EM map would still leave an extra density lobe at the C-terminus unaccounted for. This density lobe allows placement of seven additional α-helices, which nicely accommodate the TAIL-C sequences of Nup205 (Supplementary information, Fig. S12a,d). In addition, Nup188 contains a highly conserved SH3 domain in its NTD^39^ (Fig. 5a); but there is no indication of any EM density for the SH3 domain in our map. Finally, the assignment of Nup205 is supported by the length of the Tower helix in the MID domain^17^ (Supplementary information, Fig. S12a-c). Therefore, Nup205 was modeled into the density (Fig. 5b; Supplementary information, Fig. S12a).

The two molecules, named outer Nup205 and inner Nup205, interact with the short arms of the outer and inner Y complexes, respectively (Fig. 5c). The NTD of outer Nup205 directly contacts the hybrid α-helical domain of outer Nup96N/160C, whereas the tip of the Tower helix associates with the C-terminal α-helices of outer Nup85 and outer Nup160C (Fig. 5d, left panel; Supplementary information, Fig. S9a). These interactions are exactly preserved between inner Nup205 and the short arm of the inner Y complex (Fig. 5e, left panel). The TAIL of outer Nup205 interacts with outer Nup85 (Fig. 5d, left panel); but the TAIL of inner Nup205 is separated from inner Nup85 by about 30 Å (Fig. 5e, left panel). Outer Nup205 also associates with the stem of the inner Y complex from the adjacent CR subunit (Fig. 5d, right panel).

Specifically, the NTD, Tower helix, and TAIL of outer Nup205 directly contact inner Nup133, the finger helix of inner Nup107, and the N-terminal α-helical domain of inner Nup107, respectively, from the adjacent CR subunit. These interactions are unique to outer Nup205. In contrast, the TAIL of inner Nup205 associates with the Nup214 complex and the NTD contacts Nup155 that connects the inner ring (IR) to the CR^13^ (Fig. 5e, right panel).

### The Nup214 complex

The Nup214 complex is thought to contain at least three core components: Nup214, Nup88, and Nup62 (refs. 40,41), each of which has a predicted coiled-coil domain^5,28,33^ (Fig. 6a). Nup214 and Nup88 each contain a β-propeller at the N-terminus^35,42,43^. The Y-shaped EM density for the Nup214 complex is situated on the inside of the CR, with the stem projecting into the pore^12^ (Fig. 6b). The EM density for one arm allows placement of secondary structural elements and is referred to as subcomplex-1 (Sub-1) (Fig. 6c). Sub-1 contains a β-propeller that may come from Nup88, a three-stranded coiled-coil segment of 9 nm in length (referred to as CCS1), and a second coiled-coil segment of 5 nm (referred to as CCS2). Based on published information^44,45^, CCS1 and CCS2 are thought to contain the coiled-coils from the three core proteins of the Nup214 complex.

**Fig. 6.**
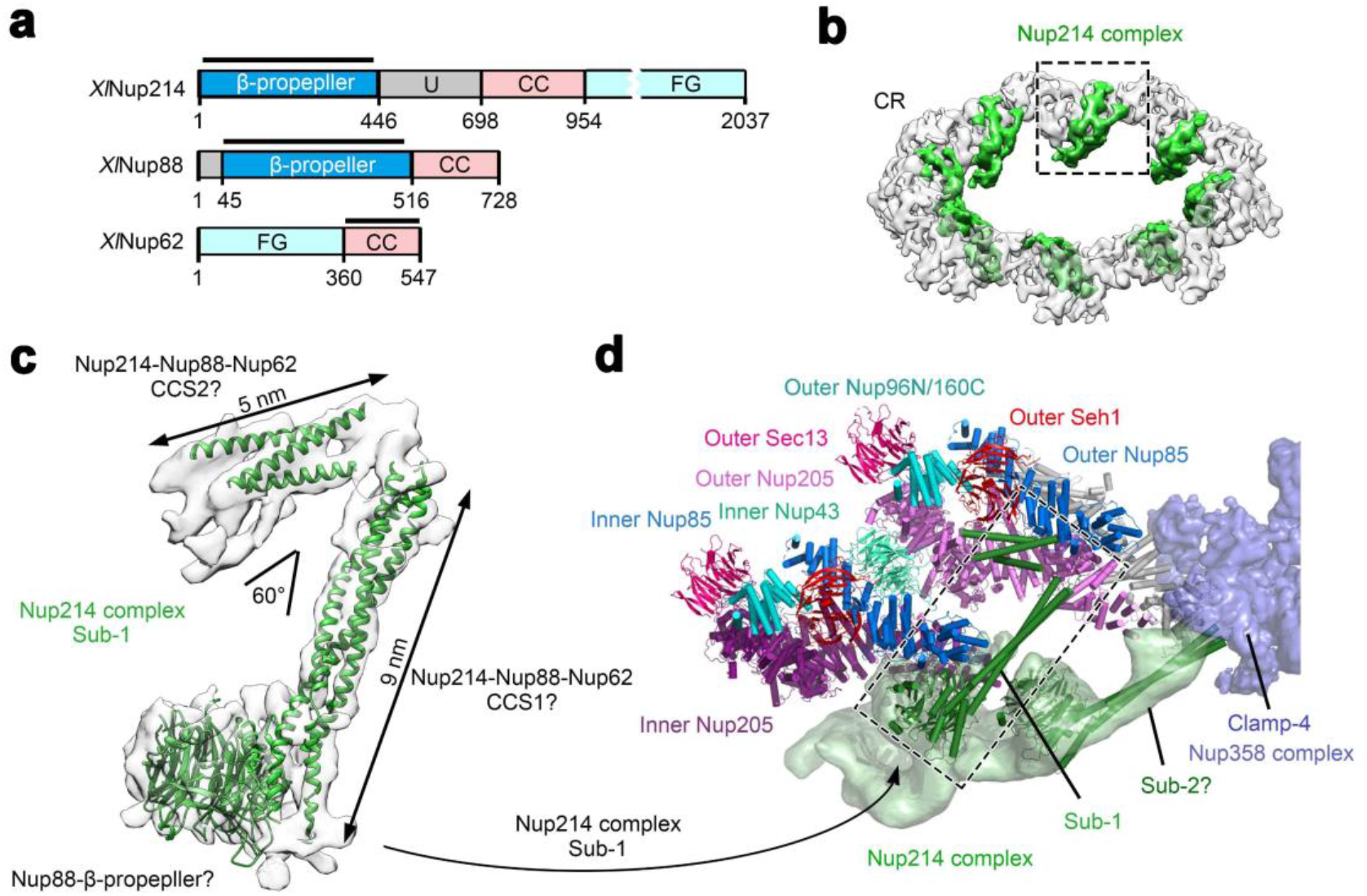
The Nup214 complex connects the outer and inner Y complexes. **a** Domain organizations of three core components of the Nup214 complex. Nup214, Nup88, and Nup62 from *X. laevis* (XlNup214, XlNup88 and XlNup62) are thought to constitute the core components of the Nup214 complex; their domain organizations are shown here. Abbreviations: CC, coiled-coil domain; U, unstructured region; FG, Phe-Gly repeats. **b** The location of the Nup214 complex in the CR. The overall reconstruction of the CR (colored by transparent grey) is shown, with that of the Nup214 complex colored green. The boxed region of the Nup214 complex is displayed in panels **c** and **d**. **c** Only a portion of the reconstruction, dubbed “sub-1” for subcomplex-1, displays fine features that allow identification of secondary structural elements. Based on published information, Sub-1 is speculated to consist of a β-propeller from Nup88 and coiled-coil segments (CCS1, CCS2) from Nup214, Nup88 and Nup62. **d** The coiled-coil domain of Sub-1 of the Nup214 complex connects the inner and outer Y complexes through direct association with inner and outer Nup85. The EM density for subcomplex-2 (Sub-2) contacts Clamp-4 of the Nup358 complex. The model of Sub-2 is highly speculative and merely shown here for reference (not included in our composite coordinates of the CR). The Nup214 complex projects into the pore of the NPC.

CCS1 of the Sub1 connects the inner and outer Y complexes through direct interactions with inner and outer Nup85 (Fig. 6d). One end of CCS1 associates with outer Nup85, whereas the other end is linked to the β-propeller of Nup88 and interacts with inner Nup85. CCS2 directly contacts the lateral side of the β-propeller from outer Seh1 and the helical ridge of outer Nup85. In contrast to Sub-1, the other arm of the Y-shaped 214 complex, referred to as subcomplex-2 (Sub-2), is characterized by weak EM density. Sub-2 connects the stem to Clamp-4 of the Nup358 complex from the adjacent CR subunit (Fig. 6d). Finally, the unassigned stem of the Nup214 complex points into the pore of the NPC (Fig. 6d).

## DISCUSSION

Cryo-ET investigations of the NPC have yielded reconstruction of the CR subunit at overall resolutions of up to 20 Å (ref. 13). The moderate resolution nonetheless allowed docking of X-ray structures of individual components and generation of a composite main-chain model for the CR subunit^13,17,23^. In this study, we report the SPA-based cryo-EM structure of the entire CR subunit at 8.0 Å resolution (Supplementary information, Fig. S3b) and the SPA-base cryo-EM reconstructions of three overlapping regions at average resolutions between 5.5 and 7.9 Å (Fig. 1 and Supplementary information, Fig. S4a-c). These reconstructions display clear features of α-helices and β-propellers, but not individual β-strands. Nonetheless, for the first time, a model of the CR has been generated with most of the secondary structural elements fitted into the EM density map (Figs. 2 & 3). Our study represents a major step forward in the structural characterization of the NPC (Supplementary information, Fig. S4d).

Nup358 functions as a cargo- and receptor-specific assembly platform^46–49^. The region that anchors Nup358 to the NPC was mapped to its N-terminal α-helical region^50^. Two density lobes previously seen in the stem and the short arm of the Y complex may come from the Nup358 or Nup214 complex^12,13^. Cryo-ET studies of human NPCs suggest the localization of Nup358 on the outer Y complex of the CR^13^. Consistent with these studies, we identified two copies of Nup358 in Clamp-1 and Clamp-2. For each of the two Y complexes within the same CR subunit, the stem is sandwiched by a pair of the Clamps, suggesting a stabilizing role. Consistent with this analysis, Nup358 knockdown led to the loss of one Y complex in the CR subunit^13^.

The Nup214 complex acts as an important platform for mRNA export and remodeling^51,52^. Nup214, Nup88 and Nup62 were suggested to form a coiled-coil subcomplex^28,44,45^. In our reconstruction, a 9-nm coiled-coil bridges a 5-nm coiled-coil and a β-propeller. This region was tentatively assigned as a core subcomplex (Sub-1) involving Nup214, Nup88 and Nup62. Quantitative mass spectrometry suggested two subcomplexes in one CR subunit in both human and yeast^15,34^. Our reconstruction suggests a potential location for a second subcomplex (Sub-2).

In our study, two S-shaped density lobes are assigned to Nup205, not Nup188 as previously proposed^12,17^. Nup205 exemplifies the remarkable versatility of a number of NPC proteins in mediating interactions with different partners (Fig. 5d,e). In addition to mediating multiple interfaces in the CR, Nup205 likely plays a role in the IR as its homologue Nup192 is also a core component of the IR^15,17,18^. Another example is Nup107. The finger helix of inner Nup107 binds the Tower helix of outer Nup205 (Fig. 5d, right panel) and the CTD of outer Nup85 from the adjacent subunit (Supplementary information, Fig. S10d), whereas the finger helix of outer Nup107 interacts with one end of the bridge domain from the Nup358 complex (Fig. 4e).

Structure determination of the NPC represents a daunting challenge for the structural biology community. Despite significant resolution improvement over previous studies (Supplementary information, Fig. S4d), our reconstruction of the CR subunit still displays little or no features for side chain and the helices mostly appear as tubular densities. As such, the directionality of the helices cannot be determined *de novo* by the reconstruction. For the Nup358 complex and a number of other proteins, the linkage among α-helices remains poorly defined by the EM map. The linkage and relationship between the bridge domain and the four Clamps are yet to be characterized. Although TAIL-C of outer Nup205 is clearly visualized, its counterpart from inner Nup205 is less well defined. This observation, together with a predicted Tower-helix in Nup188 (ref. 17) and its crosslinking to Nup85^12^, leaves the scenario not entirely impossible that the density assigned to inner Nup205 may actually represent inner Nup188. In contrast to the Nup358-containing or Nup205 region, the Nup214-contaning region has the lowest resolution. Although the EM density for the 9-nm coiled-coil region and its associated β-propeller has been assigned, we cannot rule out the possibility that it may come from other Nups.

The improved resolution was achieved through SPA, which, compared to the STA approach of cryo-ET, allows considerably higher electron dose for each movie stack. To alleviate the strong orientation bias of the NPC on the NE membrane, the movie stacks were recorded at tilting angles of 0, 30, 45, and 55 degrees^22^ (Supplementary information, Fig. S1). In particular, to ensure adequate signal-to-noise ratio, the electron dose applied was inversely proportional to the cosine value of the tilting angles. Cryo-ET studies of Human NPC have provided a starting model for this dynamic machinery.

In contrast to many other structural studies of the NPC in which a plethora of biochemical and biophysical approaches were utilized, our study relies exclusively on cryo-EM SPA. Although this aspect may represent a weakness of our study, it is our belief that this approach will ultimately result in an atomic structure of the CR, which will in turn corroborate or clarify results of the other approaches. Nonetheless, in our study, assignment of individual protein is based on a set of criteria that include previous X-ray structures, published biochemical and mass spectrometric data, sequence features and prediction of secondary structural elements. The accuracy of our model to a large extent depends on the accuracy of this information.

In contrast to vertebrate, the yeast NPC only contains one Y complex in each CR subunit^15^. Our structural observations provide a plausible explanation to this difference. First, the metazoan-specific Nup358 (refs. 5,30,31) stabilizes the CR scaffold by simultaneously interacting with the inner and outer Y complexes (Figs. 3 & 4). Second, Nup96 and Nup160 in vertebrate comprise sequences that are absent in their yeast counterparts at their N-terminus and C-terminus, respectively; Nup96N/160C directly interacts with Nup205, Sec13, Seh1 and Nup85 (Fig. 5d; Supplementary information, Fig. S9a) and may play an important role in stabilizing the Y complex scaffold. In addition, consistent with previous studies^13^, the vertebrate-specific Nup43 and Nup37 interact with Nup107 and Nup133 (Supplementary information, Fig. S10d), respectively, of the Y complex from an adjacent CR subunit to further stabilize the scaffold. Furthermore, as previously observed^17^, the marked conformational differences between inner and outer Y complexes mainly depend on the flexibility of Nup133 (Supplementary information, Fig. S11a). The β-propeller is separated from the α-helical domain by 4 nm and 2 nm in outer and inner Nup133, respectively. This variation appears to be accommodated by an extended sequence element that is uniquely present in vertebrate Nup133. Lastly, the extended finger helix of Nup107 (ref. 27) stabilizes the CR scaffold through direct interactions with Nup43, Nup85, the Tower helix of Nup205 (ref. 17), and the bridge domain of the Nup358 complex (Fig. 5d; Supplementary information, Fig. S10d). The yeast Nup84 lacks tip of the finger helix^27^ and may be incapable of bridging these interactions. In summary, our study constitutes an important framework for understanding the structure and function of the NPC.

## ACKNOWLEDGEMENTS

We thank Westlake University for the Start-up fund, the Tsinghua University Branch of China National Center for Protein Sciences (Beijing) for the cryo-EM facility and the computational facility support, and L. Zhao, X. Li, and J. Wen for technical support. This work was supported by funds from the National Natural Science Foundation of China (31930059, 81920108015, 31621092 and 31430020).

## AUTHOR CONTRIBUTIONS

X.Z. and Y.Z. prepared the sample. G.H., Y.Z., C.Z., Q.W. and J.L. collected the EM data., Y.Z., C.Z. and Q.W. processed the EM data. G.H. performed the cryo-EM SPA calculation. Q.Z., C.Y. and Q.T. provided critical advices. All authors analyzed the structure. G.H., Y.Z., C.Z. X.Z. and Y.S. wrote the manuscript. Y.S. conceived and supervised the project.

## MATERIALS AND METHODS

### Cryo-sample preparation

The protocol is similar to that as described^21^. Briefly, stage VI oocytes of *Xenopus laevis* (*X. laevis*) were sorted in the modified Barth’s saline (MBS) (10 mM HEPES, pH 7.5, 88 mM NaCl, 1 mM KCl, 0.82 mM MgSO_4_, 0.33 mM Ca(NO_3_)_2_ and 0.41 mM CaCl_2_). The nucleus was isolated and transferred into a low salt buffer (LSB) (10 mM HEPES, pH 7.5, 1 mM KCl, 0.5 mM MgCl_2_, 10 μg/ml aprotinin, 5 μg/ml leupeptin). The nuclear envelope (NE) was spread onto a freshly glow-discharged EM grid and washed using LSB before plunge-freezing. The quality of sample was examined using an FEI Tecnai Arctica microscope (Thermo Fisher Scientific) operating at 200 kV.

### Data acquisition

Grids were transferred to a Titan Krios electron microscope (FEI), operating at 300 kV and equipped with a Gatan GIF Quantum energy filter (slit width 20 eV). 23,061 micrographs were recorded with the grid tilting at angles of 0, 30, 45 and 55 degrees using a K2 Summit detector (Gatan Company) in the super-resolution mode with a nominal magnification of 64,000x, resulting in a calibrated pixel size of 1.11 Å (Supplementary information, Table S1). The total dose followed a cosine alpha scheme where the dose is inversely proportional to the cosine alpha of the tilting angle. Within each stack, the exposure time for each frame and the dose rate were kept the same. With an exposure time of 0.5 second per frame and 40 frames, the total exposure time for each 0°-tilting (tilt0) stack is 20 seconds, amounting to a total dose of 52 e^−^/Å^2^ for each micrograph. AutoEMation^53^ was used for fully automated data collection. All frames in each stack were first aligned and summed using MotionCorr^54^, with 2-fold binned to a pixel size of 2.22 Å. The output stacks from MotionCorr were further motion-corrected using MotionCor2 (ref. 55) and dose weighting was performed. The average defocus values were set between −1.0 and −3.5 μm and were estimated by Gctf^56^.

### Initial defocus estimation

To obtain reliable defocus information for the particles in the tilted micrographs, we first used Gctf to obtain the local CTF information for each particle. This initial estimation contained many outliers as visualized in 3D plot. After Gctf estimation, the tilting angle and tilting axis angle were used to generate an ideal plane that represents the grid plane. This plane was then moved up and down to find a least square fit of its position on the basis of the estimated defocus value for each particle and its X-Y coordinates. Using this fit as the starting point, a final round of gradient descent optimization using Huber loss as the loss function and the conjugate gradient method as the optimization strategy was executed to alleviate the problem of defocus estimation outliers. On the estimation of the location of the ideal plane, per particle loss was defined as the absolute deviation between Gctf-estimated defocus and predicted defocus using the ideal plane. A defocus outlier has a per particle loss that is at least twice the mean of per particle loss in this micrograph. Finally, defocus outliers were corrected for using the ideal plane.

### Initial model generation

An initial data set of 17,796 particles was used to generate an initial model. We extracted the particles using a square box of 200 pixels on each side, with a binned pixel size of 8.89 Å. To alleviate the problem of preferred orientation, we skipped the global search phase of a conventional cryo-EM data processing procedure and divided the data set into 4 subsets as defined by the tilting angles, namely the tilt0/tilt30/tilt45/tilt55 data subsets, and aligned them individually. Alignment for the individual data sets was initiated from an initial Euler angle with the tilting angle set to that of the micrograph. To avoid model bias, we used a published structure of the NPC (EMD-3103, with a reported resolution of 21.4 Å) as the initial reference and low-pass filtered to 60 Å before running the alignments. The iterative alignment was then carried out with the search range of the tilting angle restricted to be within ±3° of the result of the previous round. 100 iterations of the iterative alignment were carried out and the 4 data subsets were merged. Another 10 iterations of local search alignment with ±9° search range for each of the three Euler angles was then carried out for the merged data set. Finally, a solvent mask that only includes the cytoplasmic ring was created with a soft edge width of 13 pixels. An auto-refinement procedure using only local refinement was carried out using the created solvent mask, resulting in a reconstruction at 33 Å resolution. Two cycles of parameter refinement were then executed; each cycle has one round of auto-refinement, CTF-refinement and multi-reference 3D classification. This resulted in an improved resolution of 27 Å from 13,294 particles.

Using the 27-Å map as an initial reference and applying an initial low-pass filter of 50 Å, we performed global search K=1 3D classification, with the solvent mask covering only the cytoplasmic ring (CR). The search contained 100 iterations. The resulting data star file was directly subjected to local search auto-refinement, resulting in a reconstruction at 24 Å. Further refinement through 3D classifications, auto-refine and CTF-refine in cycles improved the resolution to 18.3 Å. Processing of the whole data set was identical to this procedure except that two rounds of multi-reference 3D classifications were executed to remove apparently bad particles and the 18.3 Å reconstruction low-pass filtered to 35 Å was used as the initial reference.

### Subunit reconstruction

We extracted subunit particles based on the alignment parameters of the 18.3 Å reconstruction. To do this, we defined a cropping center in the complex map and used Euler angles of the complex particles to deduce the 2D shift of the center of the subunit relative to the center of the complex in the micrograph. We then updated the CTF parameters using the deduced z-shift. This procedure was repeated 8 times for each complex particle to map out the location of each subunit and the corresponding Euler angles were updated accordingly. This procedure allowed us to define a 3D geometric configuration between the 8 subunits. Subunits were extracted using a box size of 200 with a binned pixel size of 4.44 Å. From this point onward, all 3D classifications and auto-refinements were carried out using only local search. An initial number of 137,795 CR subunits were extracted and resulted in an initial reconstruction at 12-Å resolution using local search auto-refinement.

### Subunit parameter refinement

Direct CTF-refinement often breaks the intrinsic geometrical restraints between subunits that belong to the same complex. To resolve this issue, we used a modified version of CTF-refinement in RELION-3.0 (ref. 57). Instead of optimizing the squared loss of each subunit, we optimized the summation of the squared loss of subunits that belong to the same complex. During the summation process, apparent outliers were down-weighted. Outliers are defined as subunits that have abnormally high or low squared loss. This practice was introduced to avoid the refinement process being biased by subunits that happened to be misaligned or covered by ice contaminants. After optimization, the defocus values of all subunits in the same complex were then updated using the same offset. On the basis of the 12-Å reconstruction obtained from 137,795 CR subunits, we performed six cycles of full parameter refinement. Each cycle comprises one round of auto-refine, modified CTF-refine and 3D classification. The final reconstruction exhibits a resolution of 9.4 Å from 59,017 particles.

### Image processing

A diagram for data processing is presented in Supplementary information, Fig. S3. The collected image stacks were motion corrected using MotionCor2 (ref. 55). From the initial data set of 23,061 micrographs, 12,399 were manually selected because they contain discernable particles (Supplementary information, Table S1). The other micrographs were not selected because they either contain few discernable particles or have ice problems. 289,306 complex particles were manually picked using RELION^58^.

Using the same procedure described in the section “Initial model generation”, we obtained a reconstruction at 17.8 Å resolution from 217,998 particles. From these particles, we extracted 1,721,077 CR subunit particles using a box size of 200 and a binned pixel size of 4.44 Å. The same procedure as the one previously described in the section “Subunit parameter refinement” resulted in a reconstruction at 8.9 Å from 1,623,386 particles. The particles were re-extracted with a box size of 256 and pixel size of 2.22 Å, resulting in a reconstruction at 8.0 Å overall resolution which was used as the initial model for further refinement of the three separate regions namely Core region, Nup358-containing region and Nup214-containing region. To improve the local EM density maps, we applied three local masks to fully cover the CR subunit.

Each of these three masked regions was subject to five cycles of parameter refinement. CTF-refinement and particle polishing were only performed on the Core region. CTF information for the Nup358-containing or Nup214-containing region was extrapolated using that of the Core region, because the Core region exhibits the highest stability and local resolution and consequently a more accurate CTF-refinement result compared to that of the Nup358-containing or Nup214-containing region. The final average resolutions for the three masked regions are: 5.5 Å for the Core region, 7.1 Å for the Nup358-containing region, and 7.9 Å for the Nup214-containing region. All reported resolutions are based on the Fourier Shell Correlation (FSC) 0.143 criterion^59^. Before visualization, all density maps were corrected for the modulation transfer function of the detector and sharpened by applying a negative B-factor that was estimated using automated procedures, effect of the solvent mask was corrected for using the high resolution noise substitution approach^60^. Local resolution variations were estimated using RELION-3.0.

### Model fitting and building

The composite coordinates of the human inner and outer Y complexes (PDB code: 5A9Q^13^) were fitted into the reconstruction of CR subunit using Chimera^61^. Atomic coordinates for the individual components of the *X. laevis* Y complex were generated through homology modeling using SWISS-MODEL^62^ and fitted into the EM density based on the coordinates of the docked human Y complexes using Coot^63^. The models were then reduced to Cα only and manually adjusted according to features of the secondary structural elements in the EM density. The disordered sequences were removed and the sequences that were absent in the initial models (e.g. several α-helices at the vertex of the Y complex) were *ab initial* modeled using Coot.

The homology model of *X. laevis* Nup205 was generated using SWISS-MODEL with *Chaetomium thermophilum* (*C. thermophilum*) Nup192 (PDB code: 5HB4 (ref. 17)) as a template. The structure of the TAIL-C domain of *X. laevis* Nup205 was manually built according to the α-helical features of the EM density. The homology model of the NTD of *X. laevis* Nup358 was generated using I-TASSER^64^ and manually adjusted to fit into the EM density of the Clamp-1 and Clamp-2 of the Nup358 complex. The poly-alanine model of a newly discovered bridge domain of the Nup358 complex was manually built based on the α-helical features of the EM density. For model building of Sub-1 of the Nup214 complex, the first coiled-coil domain of *X. laevis* Nup62-Nup58-Nup54 (PDB code: 5C3L^65^) and the β-propeller domain of *C. thermophilum* Nup82 (PDB code: 5CWW^36^) were rigidly docked into the EM density. The poly-alanine model of the CCS2 of the Nup214 complex was manually built based on the α-helical features of the EM density. The model of Sub-2 is highly speculative and not included in our composite coordinates for the CR.

After the entire composite model was generated, all proteins in the CR subunit were reduced to Cα only. Real space refinement was conducted against the 5.5-7.9 Å cryo-EM density using *real space refine program* from PHENIX^66^. Visualization and figure preparation were performed using PyMOL (www.pymol.org), Chimera^61,67^ and ChimeraX^67^.

