## Supplementary information for "Structure of the Cytoplasmic Ring of the *Xenopus laevis* Nuclear Pore Complex"

Short title: Structure of the NPC Cytoplasmic Ring

Gaoxingyu Huang^1,5^, Yanqing Zhang^2,4,5^, Xuechen Zhu^2,4,5^, Chao Zeng^3,5^, Qifan Wang^1^, Qiang Zhou^2,4^, Qinghua Tao^1^, Minhao Liu^1^, Jianlin Lei^1^, Chuangye Yan^1^, and Yigong Shi^1,2,3,4,*^

^1^Beijing Advanced Innovation Center for Structural Biology & Frontier Research Center for Biological Structure, School of Life Sciences, Tsinghua University, Beijing 100084, China

^2^Key Laboratory of Structural Biology of Zhejiang Province, School of Life Sciences, Westlake University, 18 Shilongshan Road, Hangzhou 310024, Zhejiang Province, China; ^4^Institute of Biology, Westlake Institute for Advanced Study, 18 Shilongshan Road, Hangzhou 310024, Zhejiang Province, China

^3^Tsinghua University-Peking University Joint Center for Life Sciences, School of Life Sciences, Tsinghua University, Beijing 100084, China

**SUPPLEMENTARY FIGURE LEGENDS**

**Supplementary information, Fig. S1 | Quality analysis of the cryo-EM data for the NPC from *X. laevis* oocytes.**  **a**, Representative data from the tilt-0^o^ series. A representative raw micrograph, the motion trajectory summary, and the CTF distribution are shown in the left, middle, and right panels, respectively. The results of CTF estimation using Gctf ^56^ are shown in the upper right corner of the raw micrograph (left panel). The white circles represent the estimated resolution limit. **b**, Representative data from the tilt-30^o^ series. **c**, Representative data from the tilt-45^o^ series. **d**, Representative data from the tilt-55^o^ series. As the tilting angle increases, the quality of the micrograph gradually decreases. The decrease of quality mainly originates from two aspects. First, particle motion increases as the tilting angle increases (middle panels), because tilting the sample plane projects the doming effect of the Z-direction onto the X-Y plane. Second, sample thickness is inversely proportional to the cosine of the tilting angle, resulting in an increased noise level in the micrograph. Two deliberate measures were taken to alleviate the problem of decreased quality at high tilting angles. First, the total electron dose followed an inverse-cosine scheme to increase contrast at high tilting angles. Second, more micrographs were collected at higher tilting angles. The third but unintentional measure is that a micrograph at a higher tilting angle includes more particles because the view field of on the sample plane is inversely proportional to the cosine of the tilting angle.

**Supplementary information, Fig. S2 | Structural comparison of the CR from representative studies.**  **a**, Our preliminary 18.3-Å reconstruction of the CR using single particle analysis. From top to bottom, the four panels display features of the overall CR (colored marine), Nup205 or Nup188 (magenta), the Nup214 complex (green) and the Nup358 complex (purple). **b**, Cryo-ET structure of the *X. laevis* NPC using sub-tomogram averaging (STA) (EMD 3005-3008)^14^. **c**, Cryo-ET structure of the *Homo sapiens* (*H. sapiens*) NPC using STA (EMD-3103)^13^. Although the EM map of human NPC was used as the initial reference, our reconstruction more closely resembles the cryo-ET reconstruction of the *X. laevis* NPC in terms of both overall shape and fine features. In particular, the Nup214 complex resembles the cryo-ET *X. laevis* NPC more than the cryo-ET human NPC. Therefore, our reconstruction was not biased by the initial reference used. Subsequent analysis at the CR subunit level shows a plethora of structural details that were previously unrecognized in low-resolution reconstructions.

**Supplementary information, Fig. S3 | Cryo-EM data processing.**  **a**, A flowchart diagram of preliminary cryo-EM data processing. Only 17,796 particles from 860 movie stacks were used. Details of the data processing are described in the Materials and Methods. **b**, A flowchart diagram of the complete cryo-EM data processing. The CR subunit is divided into three overlapping regions: the Core region, the Nup358-containing region, and the Nup214-containing region. The final reconstructions of the Core region, the Nup358-containing region, and the Nup214-containing region display average resolutions of 5.5 Å, 7.1 Å and 7.9 Å, respectively.

**Supplementary information, Fig. S4 | The quality of cryo-EM reconstruction for the CR subunit.**  **a**, The quality of cryo-EM reconstruction for the Core region. The EM density, angular distribution of the particles used for the final reconstruction, and directional FSC of the Core region are shown in the left, middle, and right panels, respectively. In the angular distribution plot, each cylinder represents one view and the height of the cylinder is proportional to the number of particles for that view. **b**, The quality of cryo-EM reconstruction for the Nup358-containing region. **c**, The quality of cryo-EM reconstruction for the Nup214-containing region. All directional FSC curves were prepared using the following website: <https://3dfsc.salk.edu>^22^. **d**, Comparison of our cryo-EM reconstruction of the *X. laevis* CR with that of the *H. sapiens* CR^13^. The same regions of the reconstruction are shown for the cryo-EM reconstruction of the *X. laevis* CR (upper panels) and the cryo-ET reconstruction of the *H. sapiens* CR^13^ (lower panels).

**Supplementary information, Fig. S5 | Representative EM density maps for Nup85, Seh1 and Nup43 of the Core domain.**  **a**, The overall EM density map of Nup85 and Seh1. The EM density maps for Nup85 and Seh1 from the inner and outer Y complexes are shown in the left and right panels, respectively. **b**, Representative EM density maps for a number of discrete α-helices from inner Nup85. The EM density maps for four pairs of HEAT repeats are shown in the upper panels. **c**, The EM density maps for the β-propeller domains of inner Seh1 (left panel) and inner and outer Nup43 (right panels). One blade in the Seh1 β-propeller comes from Nup85. All EM density maps in this figure were prepared using the masked Core region map with a contour level between 15σ and 25σ.

**Supplementary information, Fig. S6 | Representative EM density maps for Nup96N/160C and Sec13 of the Core domain.**  **a**, The EM density map of Nup96N/160C. The EM density maps for the NTD of Nup96 and the CTD of Nup160 are shown in the upper panels. Representative EM density maps are shown in the lower panels for selected α-helices of the inner and outer Nup96N/160C. **b**, The EM density maps of the β-propeller domains in the inner (left panel) and outer (right panel) Sec13. Each β-propeller only has six blades. The unoccupied density for one blade in both cases comes from Nup96. All EM density maps in this figure were prepared using the masked Core region map with a contour level between 15σ and 25σ.

**Supplementary information, Fig. S7 | The EM density maps for the Nup358 complex.**  **a**, The EM density maps for the bridge domain (left panel) and selected α-helices from the bridge domain (right panel). **b**, Representative EM density maps for selected regions of the Nup358 complex. Shown here are the EM maps for the N-terminal helices of Clamp-1 and Clamp-2 and the U-domain from Clamp-1. All EM density maps in this figure were prepared using the masked Nup358-containing region map with a contour level between 15σ and 25σ.

**Supplementary information, Fig. S8 | The EM density maps for Nup205.** **a**, The overall EM density maps for the outer and inner Nup205. **b**, The EM density maps for the MID domain of outer Nup205, which contains a characteristic Tower helix. The ARM repeat and HEAT repeat exhibit contrasting features for their EM density. **c**, Representative EM density maps for a number of discrete α-helices of outer Nup205. All EM density maps in this figure were prepared using the masked Core region map with a contour level between 15σ and 25σ.

**Supplementary information, Fig. S9 | The EM density maps for representative interfaces in the CR subunit.** **a**, Representative EM density maps for proteins that interact with Nup205. Shown here are the local EM density maps for the proteins that interact with the NTD (left panel), the Tower helix (middle panel) and TAIL (right panel) of outer Nup205. The individual protein components and their associated density maps are color-coded. **b**, The EM density maps surrounding the N-terminal domain of outer Nup96. All EM density maps in this figure were prepared using the masked Core region map with a contour level between 15σ and 25σ.

**Supplementary information, Fig. S10 | Interactions among the Y complexes.** **a**, The final coordinates of the inner and outer Y complexes are placed into the EM density map. Two views are shown. **b**, Interactions among the Y complexes within the same CR subunit and between two adjacent subunits. The inner and outer Y complexes interact with each other within the same subunit. A pair of Y complex from one subunit associate with its counterpart from the adjacent subunit in a head-to-tail fashion. **c**, A close-up view on the interface between the inner and outer Y complexes within the same subunit. **d**, A close-up view on the interface between two neighboring CR subunits. Five areas of interactions are indicated by dashed oval circles.

**Supplementary information, Fig. S11 | Structural comparison among the Y complexes.**  **a**, Structural comparison between the inner and outer Y complexes within the same CR subunit. Two perpendicular views are shown. The inner and outer Y complexes are colored salmon and yellow, respectively. **b**, Structural comparison between the *X. laevis* Y complex and the *H. sapiens* Y complex. Coordinates of the inner and outer Y complexes from *X. laevis* are superimposed with those from *H. sapiens*^13^. The overall conformation and main features of the Y complexes remain very similar between these two species. The only notable difference is that the top and bottom faces of the Nup43 β-propeller in the *X. laevis* Y complexes are opposite of those in the human Y complexes. **c**, Structural comparison between the *X. laevis* Y complex with the *Myceliophthora thermophila* (*M. thermophila*) Y complex^23^. **d**, Structural comparison between the *X. laevis* Y complex with the *Saccharomyces cerevisiae* (*S. cerevisiae*) Y complex^19^. The overall conformation of the *X. laevis* Y complex is similar to that of the *S. cerevisiae* Y complex, but numerous local structural variations are present. There are no *S. cerevisiae* homologues for the *X. laevis* Nup43 and Nup37.

**Supplementary information, Fig. S12 | The basis for the assignment of Nup205.**  **a**, The overall EM density map is more consistent with that for Nup205. The presence of continuous EM density for additional α-helices beyond the predicted C-terminal end of Nup188 favors the assignment of Nup205. Additionally, the length of the EM density for the Tower helix is consistent with that of CtNup192 or *X. laevis* Nup205. **b**, Comparison of the Tower helices between *X. laevis* Nup205 and CtNup192. Inner and outer Nup205 is compared to CtNup192 in the left and right panels, respectively. **c**, The length of the predicted Tower helix in Nup205 is similar to that of CtNup192 (PDB code: 5HB4)^17^. Shown here is a sequence alignment of the Tower helix region between CtNup192 and Nup205 of three vertebrate species^33,68^. **d**, The EM density that is connected to the rest of the EM density for Nup205 or Nup188. Seven α-helices (two of them are longer than the other five and are slightly bent in the middle) can be accommodated by the EM density map. Based on the location and continuity of the EM density, these seven α-helices are assigned to the C-terminus of Nup205 (TAIL-C).

**Supplementary information, Table S1 | Statistics of cryo-EM data collection and analysis.**

**Supplementary information, Table S2 | Summary of model building for the CR subunit of *Xenopus laevis* NPC.**
